# Urine cf-Nucleosomes: A Non-invasive Window into Human Physiology and Disease

**DOI:** 10.1101/2024.09.01.610671

**Authors:** Matan Lotem, Israa Sharkia, Batia Azria, Tal Falik-Michaeli, Nir Friedman

## Abstract

Urine contains fragments of cell-free DNA, which hold valuable molecular insights into the processes occurring within the urinary system and the whole body. It is unknown whether these fragments are in the form of chromatin, as in the cell of origin, and whether they maintain the cell of origin chromatin modifications. Here, we examine these questions using cell-free chromatin immunoprecipitation followed by sequencing (cfChIP-seq) on human urine. We demonstrate that we can capture cell-free nucleosomes (cf-nucleosomes) from urine and that these preserve multiple histone post-translational modifications indicative of activation and repression. By analyzing these modifications, we identified the primary tissues contributing to cf-nucleosomes in a cohort of healthy individuals. Notably, we observe distinct populations of circulating cf-nucleosomes in urine samples from healthy donors with a contribution from the kidney, which are not detected in matched urine exfoliated cells or matching plasma samples. We further show that, at most, a negligible amount of urine cf-nucleosomes originates from plasma, suggesting that kidney filtration excludes plasma-circulating nucleosomes from urine. Additionally, we show that urine cf nucleosomes can report pathologically driven changes in the urine of bladder cancer patients, reflecting tumor-associated transcriptional programs and immune responses. Our findings highlight the potential of urine cf nucleosomes as accessible, noninvasive biomarkers for both basic research in renal physiology and monitoring urinary pathologies.

**Key Findings:**

- Urine cell-free nucleosomes exist and retain multiple histone marks that are informative of gene promoters and enhancers.
- Urine cfChIP-seq identifies bladder, kidney, and immune cells as the major contributing organs to the pool of urine cell-free nucleosomes.
- The populations of cell-free nucleosomes in urine and blood are distinct and primarily disjoint, suggesting that few, if any, nucleosomes cross the blood-urine interface.
- Urine cell-free nucleosomes reflect pathologically driven changes in tumors and immune cells responding to the tumor.

## Graphical abstract

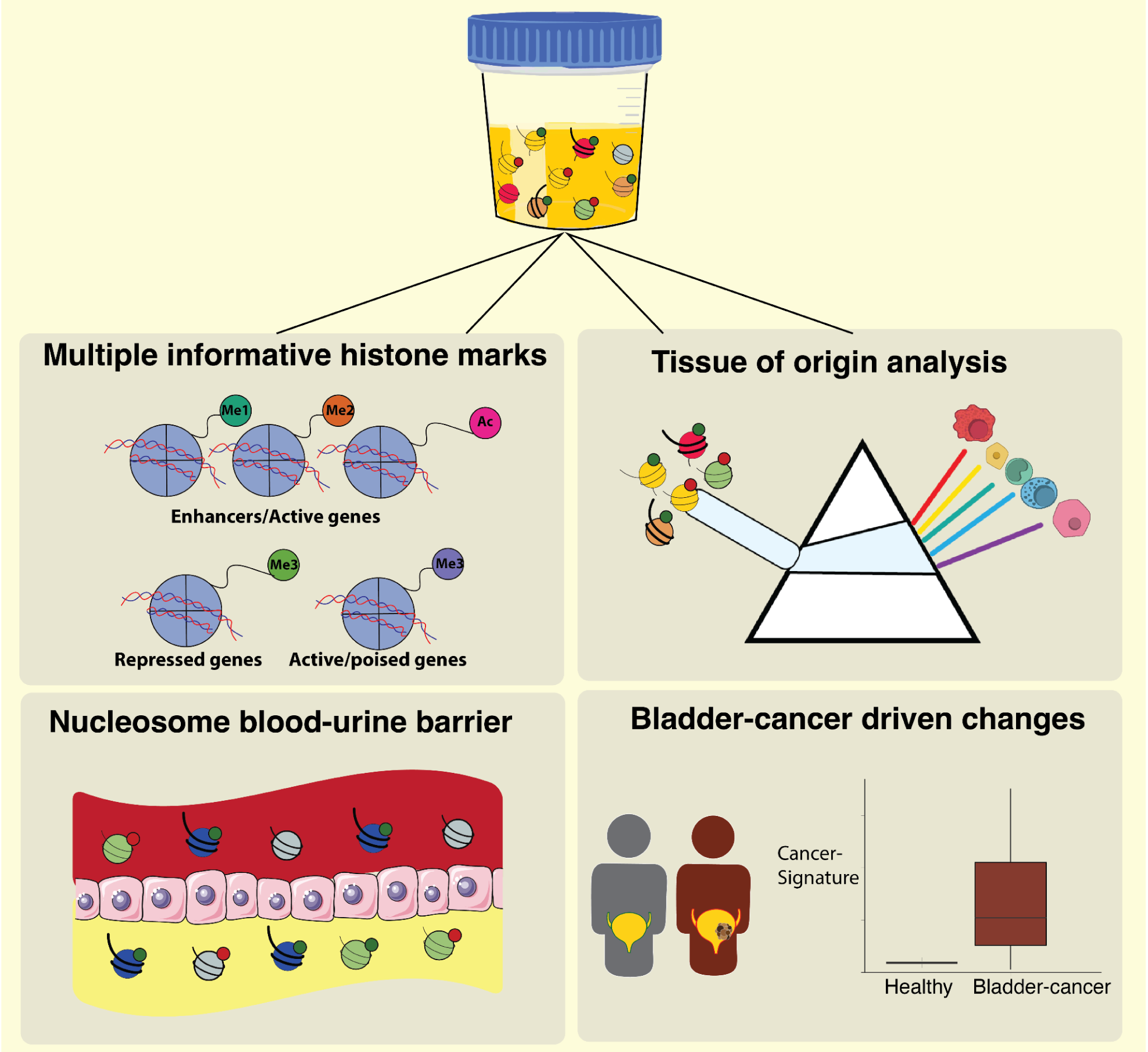

## Introduction

A long-standing goal in diagnostic medicine is noninvasively monitoring an individual’s physiological status. Noninvasive diagnostic tools are essential in the early stages of the diagnostic process when the clinician has to consider multiple hypotheses. Blood and urine are two rich sources of information. Each collects molecules from large parts of the human body. Correspondingly, there is a wide range of blood and urine diagnostic tests for metabolites, proteins, nucleic acids, and more.

Recently, nucleic acids in urine have gathered interest as potent biomarkers^1^. cfDNA from urine has proven to be a source of biomarkers for various pathologies, including urogenital cancers such as prostate and bladder cancer^2^, but also non-urogenital cancers such as glioma^3^ and colorectal cancer ^4^. Urinary cfDNA has also shown potential applications in monitoring urinary tract infections and organ transplant rejections ^5,6^ and in prenatal diagnosis ^7^. Notably, urinary cfDNA can originate from healthy and diseased tissues, making it a valuable biomarker for assessing an individual’s physiological and pathological states ^8^.

Urinary cfDNA is reported to degrade within minutes to less than 3 hours by active DNase I in the urinary tract ^9–12^. Thus, it originates from recent, close-to-urination time events. Urinary cfDNA originates from two sources. The first is from cells along the urogenital tract that shed DNA fragments through multiple pathways ^2^. The second is from transrenal DNA fragments that pass from blood circulation, traversing kidney filtration ^13^.

Some cell death pathways (e.g., apoptosis) result in the release of short chromatin fragments (complexes of DNA and proteins). The basic repeating units of chromatin are nucleosomes, each consisting of approximately 147 base pairs of DNA wrapped around a histone protein core composed of an octamer of histones H2A, H2B, H3, and H4 ^14^. During programmed cell death, DNAses cut accessible linker regions between nucleosomes, and chromatin fragmentation displays a ladder of mono-, di-, and tri-nucleosome-length DNA fragments reflecting the underlying chromatin organization^15^.

Multiple lines of evidence show that most of the circulating cfDNA in the blood originates from fragments of nucleosome origin and that in the circulation, most of cfDNA is still in intact complexes with histones ^16–18^. Analogous questions arise in the context of urine cfDNA: Is it of nucleosomal origin and to what extent is it maintained as intact chromatin fragments? In particular, is transrenal cfDNA in the form of intact chromatin fragments?

Histone proteins have been described in urine ^19^, however it is not known if they represent intact nucleosomes. Necrotic and apoptotic cells release chromatin into their immediate environment ^20^. Thus, we expect to find cell-free nucleosomes from dying urogenital cells in urine. Conversely, the process of urine formation by glomerular blood filtration entails that large proteins with molecular weight >45 kilodaltons rarely pass filtration ^21–23^. Thus, a healthy kidney is expected to exclude plasma circulating nucleosomes (∼179 kilodaltons) ^24^ from the urine.

The harsh urine environment with active DNAse digestion further complicates the answer to these questions. The length distribution of urine cfDNA suggests that most cfDNA fragments are partially digested ^9,12^, in contrast to plasma cfDNA, which has characteristic peaks at mono- and di-nucleosome lengths (∼170bp and 320bp).

In living cells, nucleosomes serve as a dynamic platform for DNA-associated processes. These processes involve chromatin-modifying enzymes that deposit and remove post-translational modifications of histones, such as methylation, phosphorylation, and acetylation, and chromatin-binding proteins that recognize specific modifications^14^. These chromatin modifications play central roles in transcription, replication, and DNA repair ^25–27^. Histone marks offer clear and well-defined relationships to cellular processes, making chromatin analysis valuable for understanding cell identity and state ^28–30^. For example, cardiomyocytes express the TNNT2 gene, and so only in this cell type would we encounter TNNT2-promoter nucleosomes marked with expression-corresponding marks (e.g. acetylations, phosphorylation, tri-methylation, etc). Some cancers, such as mixed lineage leukemia^31^, are characterized by epigenetic misregulation resulting in aberrant histone-markings of oncogenes - indicative of cancerous cell-state ^32,33^.

Recent works ^34–36^ show that many histone modifications survive in plasma cell-free nucleosomes. By assaying them, we recover rich information about gene activation, regulatory processes, and more. Capturing chromatin fragments with a specific modification (e.g., one associated with active promoters) and sequencing the associated DNA, we can identify genomic locations that had the modification in the cell of origin (e.g., active promoters in these cells) and learn about their identity and state.

These results raise the question of whether urine nucleosomes, despite the harsh conditions, also carry cell-of-origin histone modifications and whether assaying these can provide insights about their origin.

To answer these questions, we performed, for the first time, cell-free chromatin immunoprecipitation and sequencing (cfChIP-Seq ^36^) in human urine. We targeted both active (H3K4me1/2/3, H3K27ac) and repressive (H3K27me3) histone marks and showed that urine contains cell-free nucleosomes that retain multiple histone marks. Using cfChIP-seq against H3K4me3 (an active/poised promoter histone mark) in a cohort of healthy samples, we characterize the cell of origin of urine cell-free nucleosomes. Moreover, cfChIP-seq profiles of a cohort of samples from non-muscle invasive bladder cancer patients (NMIBC) highlight pathology-driven changes. By comparing matched urine and plasma samples in healthy individuals as well as women pregnant with male fetuses, we show that negligible amounts of nucleosomes pass from plasma to urine, suggesting that urine and plasma are non-overlapping sources of epigenetic information. These results highlight the information urine cell-free nucleosomes can provide on the molecular states of cells in the renal and urinary systems, making them valuable biomarkers for liquid biopsy.

## Results

### Modified circulating nucleosomes in human urine

We performed urine cfChIP-seq on urine samples from healthy individuals with antibodies targeting marks of accessible/active promoters (H3K4me3 n=51, H3K27Ac n=7), enhancers (H3K4me2 n=13, H3K4me1 n=13, H3K27Ac n=7) and repressive heterochromatin (H3K27me3 n=5) (Methods, Fig. 1A-C, S1A). Assays were performed on 2 ml of urine supernatant and yielded sequencing libraries with an average complexity of >13 million unique reads (Methods) and an average of >80% alignment rates (Supplementary Table 1). These quantities are consistent within an order of magnitude within total cfDNA concentrations in urine (Supplementary Note 1), suggesting that a non-trivial fraction of cfDNA in urine is nucleosomal.

**Figure 1.**
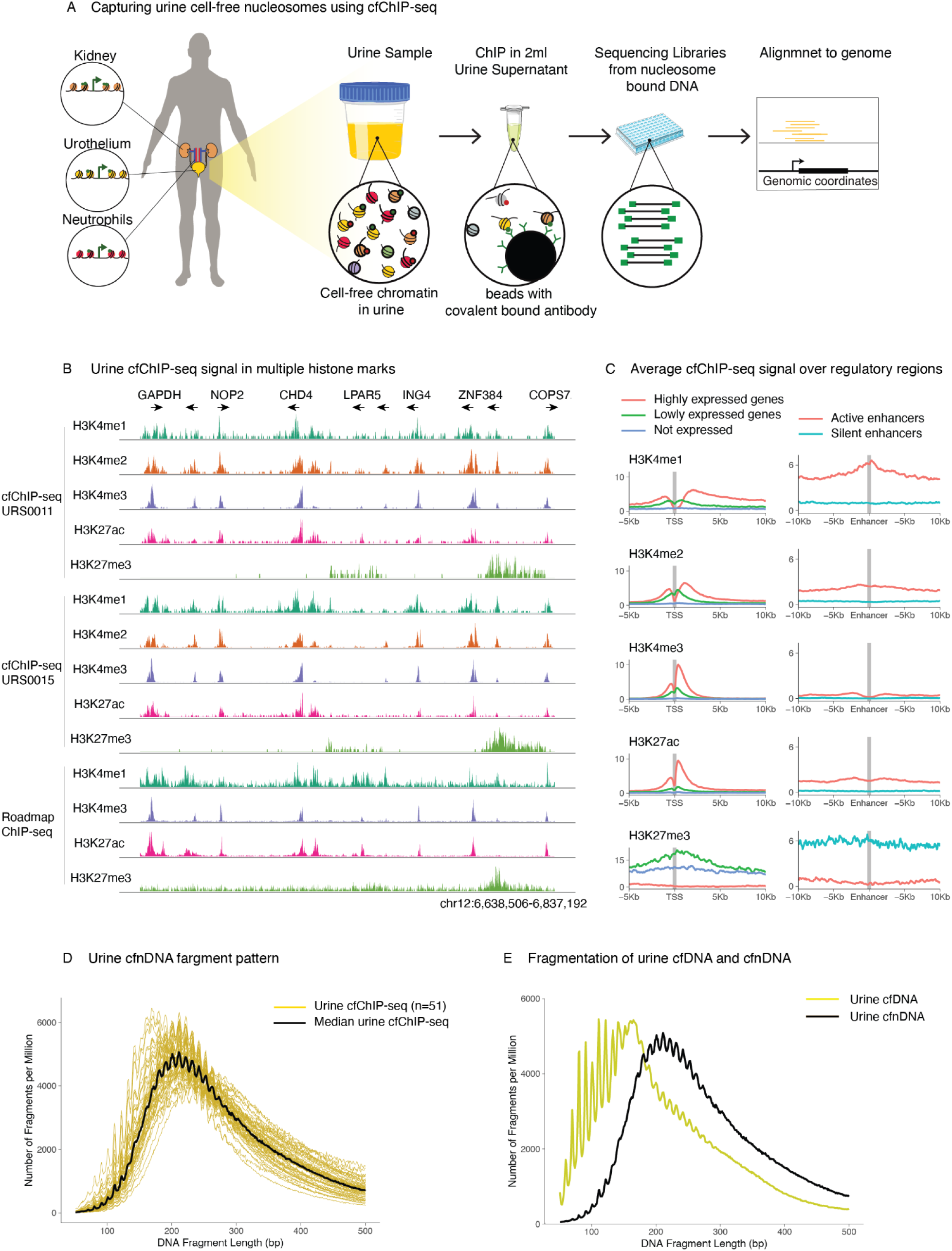
cfChIP-seq of multiple histone modifications in human urine. A. Urine cfChIP-seq outline. Different cells release chromatin fragments into the urine. These fragments are immunoprecipitated using beads with covalently bound antibodies. DNA fragments are released and sequencing libraries are generated. B. Genome-browser view of histone modifications signal. Three sets of tracks are shown: Two from cfChIP-seq of healthy donors and one from published epithelial-tissue ChIP-seq data ^44^. C. Metagene of normalized cfChIP-seq signal (Methods) over promoters and enhancers. Line colors indicate activity levels of the genes based on >2000 genes and enhancers from published epithelial-tissue ChIP-seq data^44^. Graphs scale are coverage of fragments per million. D. cfChIP-seq DNA fragment size distribution. Thin yellow lines show DNA fragments from 51 urine cfChIP-seq samples of various histone modifications (H3K27ac n=6, H3K27me3 n=5, H3K4me1 n=13, H3K4me2 n=13, H3K4me3 n=51), black line shows the median of all samples. E. Fragmentation patterns of urine ChIP-seq cfnDNA and urine cfDNA. Line shows a median of n=51 repeats in cfnDNA (black) and n=10 repeats in cfDNA (yellow).

### Circulating nucleosome modifications originate from intact cells

Mapping the DNA sequences enriched in immunoprecipitated circulating nucleosomes to the genome, we observe the show stereotypical patterns in promoters and enhancers (Fig. 1B-C, S1A-B) for both active marks (H3K4me1/2/3, H3K27ac) and repressive marks (H3K27me3). These patterns match the profiles observed in tissue ChIP-seq (Fig. 1B-C, S1B). And as in tissue ChIP-seq, we observe H3K4me3 at promoters of active and poised genes, H3K4me2/1 at regulatory regions (promoters and enhancers), H3K27ac at active regulatory regions, and H3K27me3 surrounding repressed regulatory regions.

These results show that marks showed the expected enrichment observed in intact cells. Given the high specificity of histone methyltransferase enzymes and their relatively low abundance in cells^37^, these marks are highly unlikely to be deposited after cell death, suggesting that histone modifications on circulating nucleosomes originate from intact cells.

### Circulating nucleosomes are a distinct subpopulation of cfDNA

ChIP enriches for histone-bound DNA, and thus we will refer to the products of ChIP cell-free nucleosomal DNA (cfnDNA). The lengths of cfnDNA fragments captured as a continuum in the range of mono-nucleosome (∼170 bp) to tri-nucleosome (∼400 bp) fragments (Fig. 1D-E, S1C-D). In contrast, urine cfDNA fragment lengths span a broader range from short (< 100 bp) to long (>1000 bp) ^8,38^. Specifically, the cfDNA pool includes many short fragments (<100bp) that do not appear in the cfnDNA pool. This discrepancy is not due to technical sources (library preparation and sequencing protocols): we sequenced urine cfDNA using the same protocols used for cfChIP-seq (Fig 1E) and reproduced the same discrepancy in the short fragments distribution. These results suggest that cfChIP-seq captures nucleosomal DNA fragments, which by nature are longer than 100bp. In contrast, cfDNA sequencing captures an additional pool of non-nucleosomal fragments, including fragments shorter than 100bp.

Both urine cfDNA and cfnDNA have a 10 bp length periodicity, consistent with DNaseI digestion pattern (Fig 1E, S1D) ^3,39,40^. Together with the fragment length distribution, this suggests that the fragments recovered by ChIP-seq undergo multi-phase digestion where nucleosomal ladder fragments (presumably digested at cell death) are subsequently digested in the urine by DNaseI. This is consistent with the high abundance of DNaseI in urine, which is also more active in urine than in plasma ^41,42^, and partial unwrapping of nucleosomes mainly due to the high salt and urea concentrations in urine ^43^.

### Tissues of origin of the urine cf-nucleosome pool

To further analyze the tissue origin of urine cf-nucleosomes, we focused on the H3K4me3 histone modification, which marks active or poised promoters and was shown to correlate with gene transcription and be informative of cell identity and cell state ^36^. We used published ChIP-seq data to define tissue-specific signatures as promoters with high signal only in one tissue-type (Methods, Fig. 2A-B).

**Figure 2:**
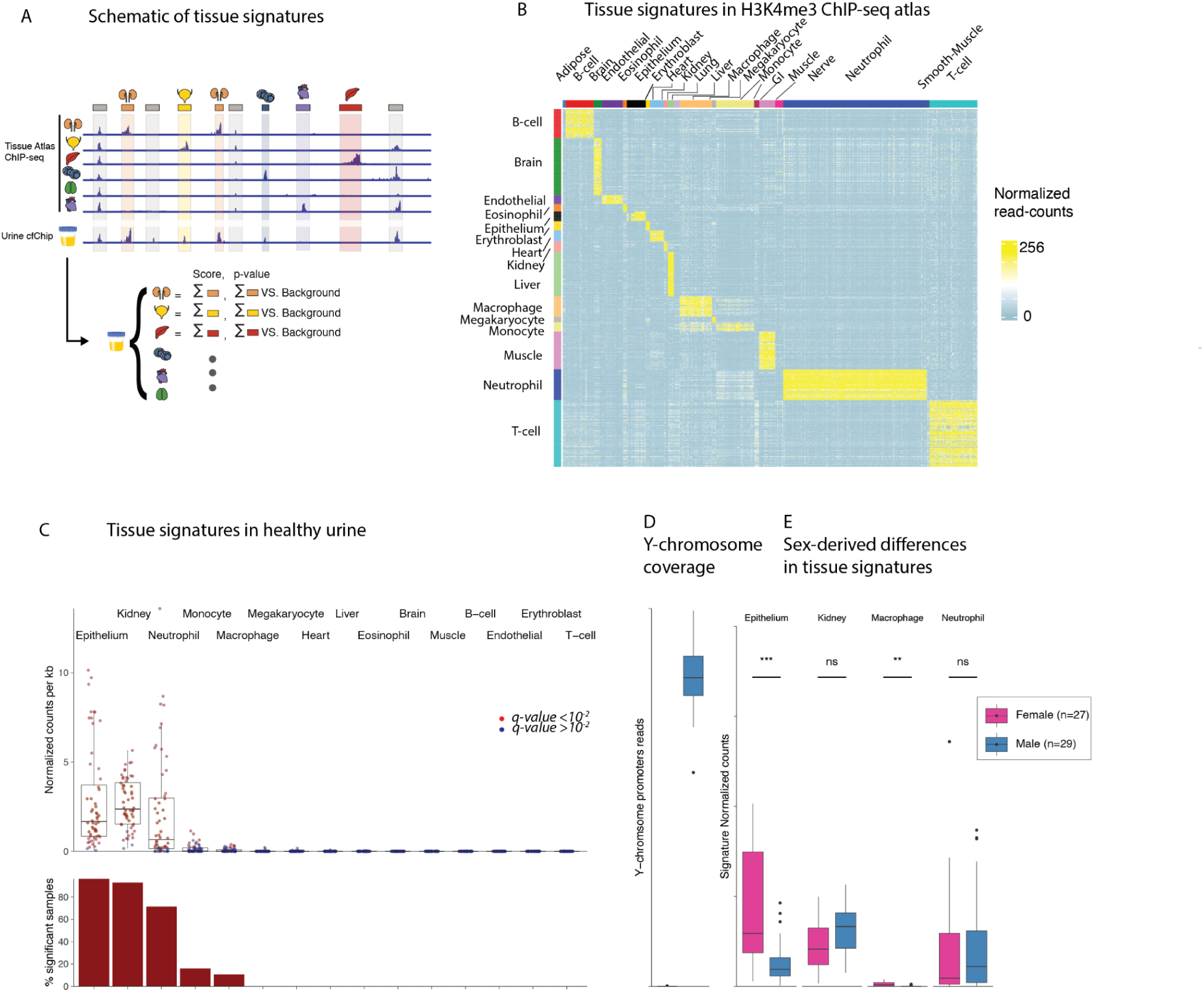
cfChIP-seq detects cfDNA cell of origin. A. Schematic of genomic regions used as tissue-specific signatures based on reference ChIP-seq tissue atlas. For each tissue (rows in genome browser view) we identify genomic locations that have high signal only in the target cell type (column with colored matching icons. Colored squares represent tissue specific windows. Gray windows are non-specific or background windows). Given a urine cfChIP-seq sample (bottom row and right-side of illustration), we sum the signal at signature locations and test against the null hypothesis of non-specific background coverage (Methods). B. Heatmap of tissue signatures (y-axis) in H3K4me3 chip-seq signal of different tissues in the tissue-atlas (x-axis). Color indicates the normalized read-count of each genomic-window in each tissue. C. Top, Tissue signatures in cfChIP-seq in 56healthy samples from 21 donors. Each dot is a sample. Box limits: 25–75% quantiles, middle: median, upper (lower) whisker to the largest (smallest) value no further than 1.5× interquartile range from the hinge. Dots marked in red indicate values significantly higher than background levels (Methods). Bottom, percent of samples with significant signal for each signature. D. Normalized promoter coverage in the Y-chromosome in urine samples from female (27 samples from 9 donors) and male (29 samples from 12 donors) donors. E. Tissue signatures in female and male donors as in C. Box plots description as in B. Wilcox-test p-values for comparison between the sexes are presented for each tissue signature. */**/***/**** correspond to *P* < 0.05/0.01/0.001/0.0001.

In nearly all healthy urine donors, we observed a strong signal of urinary epithelium, kidney and neutrophils, and in a few samples, macrophages, monocytes, B-cells, and T-cells signatures. We do not see contributions from other tissue types tested (Fig. 2B, S2A). These are the expected cell types according to observations from cfDNA methylation in urine ^9^ and single-cell RNA-seq of urine-exfoliated cells ^45,46^.

Men and women differ genomically and anatomically, which are reflected in assay results. We did not observe sex differences in assay quality (Supplementary Figure S2C). Yet, urine cfChIP-seq profiles differ between the sexes. Beyond the expected differences in sex chromosomes read coverage (Fig. 2C), a comparison of urine tissue signatures finds several sex-based differences (Fig. 2D, S2B). We observed significantly more urothelium in females compared to males. This is concordant with previous observations ^46,47^, and with increased cell exfoliation in female urethra serving as a defense mechanism against pathogens ^48^. We observed higher macrophage levels in females but did not observe significant sex-based differences in the levels of kidney and neutrophils.

### Urine cf-nucleosomes do not originate solely from exfoliated cells

Cells shed throughout the urinary tract reach the urine. A simple model would suggest that all cfnDNA originates from these exfoliated cells. As reported previously ^46,47^, male urine contains negligible amounts of exfoliated cells, while female urine contains thousands to tens-of thousands of exfoliated cells. According to the simple model we would expect the cell-type composition of cfnDNA to mimic the population of exfoliated cells.

To test this hypothesis, we compared urine cf-nucleosomes to chromatin of urine-derived exfoliated cells from the same individual (Fig. 3A-B, S3A) and systematically across multiple individuals (Fig. 3C).

**Figure 3:**
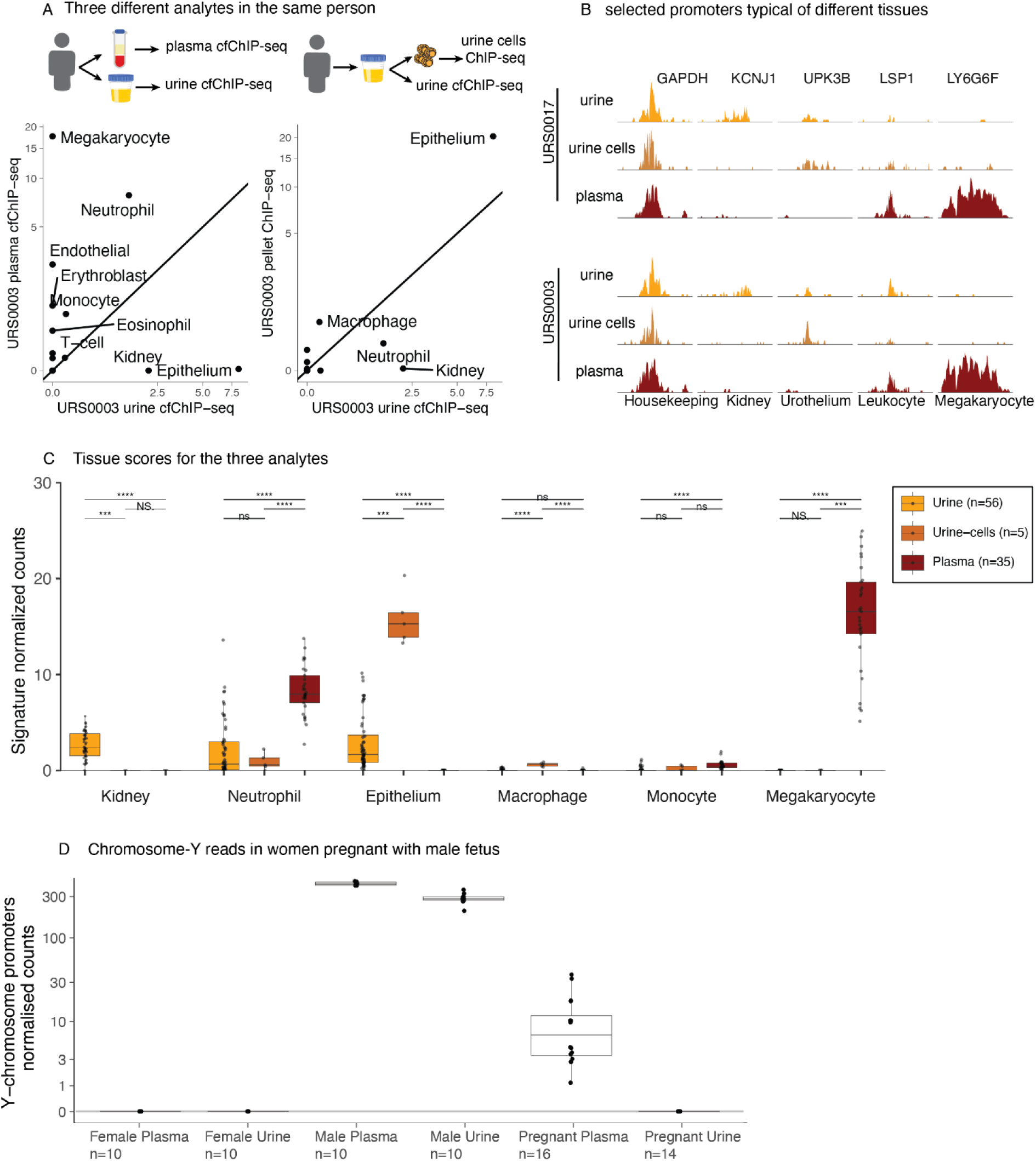
Differences in promoter signal of three analytes. A. Comparison of urine cfChIP-seq (x-axis) vs. donor matching plasma cfChIP-seq (y-axis left) and ChIP-seq of urine-cells (y-axis right). Dots represent signature-scores of labeled tissues. B. Genome browser comparing the three analytes (urine cfChIP-seq, plasma cfChIP-seq and urine-cells ChIP-seq) in two individuals. Each column features a promoter representative of a tissue-type. C. Tissue signature distribution of multiple samples in the three analytes: urine cfChIP-seq (56 samples from 21 donors), urine-cellsChIP-seq (5 samples from 4 donors) and plasma cfChIP-seq (35 samples from 35 donors). Boxplot description and statistical test as in figure 2D. D. Normalized promoter coverage in the Y-chromosome in urine and plasma samples from healthy non-pregnant females, males and females pregnant with male fetuses.

In both urine cfChIP-seq and exfoliated cells ChIP-seq, we observe a significant signal associated with the epithelial signature. This aligns with previous findings that the majority of exfoliated cells in urine are of epithelial origin ^46,49^. Consistent with earlier reports indicating that renal cells are rare in healthy urine samples ^46^, we did not detect kidney-derived nucleosomes in exfoliated cells. However, we detect kidney-derived nucleosomes in cfnDNA (Fig. 3C, Fig. S3A). This suggests that, in healthy donors, most kidney-derived cfDNA arises from *in-situ* cell death rather than exfoliation and subsequent cell death within the bladder.

### Glomerular filtration blocks plasma cf-nucleosomes from reaching the urine

In plasma, we observe megakaryocyte-derived and monocyte cfnDNA due to megakaryocyte cell death in the bone marrow and monocyte turnover in the circulation ^50,51^. Despite a large contribution in all plasma samples, we do not observe megakaryocyte signatures in both urine fractions, suggesting that in healthy donors, these nucleosomes are effectively filtered in the kidney.

For a more definite test, we collected matched urine and plasma samples from pregnant women ranging from gestational ages weeks 10 up to 40. In 16 samples from 6 women carrying a male fetus, we observe reads from Y chromosome promoters in H3K4me3 from plasma. However, we do not observe any reads from the Y chromosome in urine samples from the same women. We see an increase in chromosome Y signal as gestational age increases in plasma but not in urine (Fig. S3C). Fetal cfDNA in general, and specifically from chromosome-Y origins can be captured in urine cfDNA ^7,52,53^. Taken together, these results show that cf-nucleosome bound cfDNA in the plasma does not reach the urine under healthy conditions.

### Urine in cell-free chromatin changes in bladder cancer patients

To test the utility of urine cf-nucleosomes in differentiating between transcription patterns within the source tissue, we performed the cfChIP-seq on urine samples from patients diagnosed with non-muscle invasive bladder cancer (NMIBC, Supplementary Table S2). This cancer originates from the urothelium, which lines the inner surface of the bladder. Bladder cancer is one of the most common cancers in humans, with more than half a million new cases worldwide and hundreds of thousands of subsequent deaths ^54^. Bladder cancer affects ∼4 times more men than women, and disease incidence increases with age. Over 70% of patients diagnosed with bladder cancer are diagnosed with NMIBC ^55^. NMIBC has a high probability of recurrence and progression ^56^. Currently, the gold-standard in NMIBC monitoring is frequent cystoscopies ^57^, which adversely affect the patients’ quality of life ^58^. Non-invasive approaches to monitoring and studying this cancer are an unmet need.

H3K4me3 urine cfChIP-seq performed on samples from 25 male NMIBC patients shows hundreds of gene promoters with increased cfChIP-seq signal compared to a healthy reference (Fig. 4A). To systematically evaluate these differences, we compared these NMIBC urine samples to urine samples from a cohort of 29 self-reported healthy males (Supplementary Table S2). At the tissue-signature level, the NMIBC-cohort samples show an increase in epithelium and immune cells compared to the healthy cohort (Supplementary Fig. S4A) presumably driven by the cancer’s contribution to the cf-nucleosome pool. However, to better understand this contribution, we tested for differentially marked genes between the two cohorts and identified 177 genes with significantly elevated coverage and 61 genes with considerably decreased coverage (Fig 4B; Methods), indicating changes in gene regulation between the groups.

**Figure 4:**
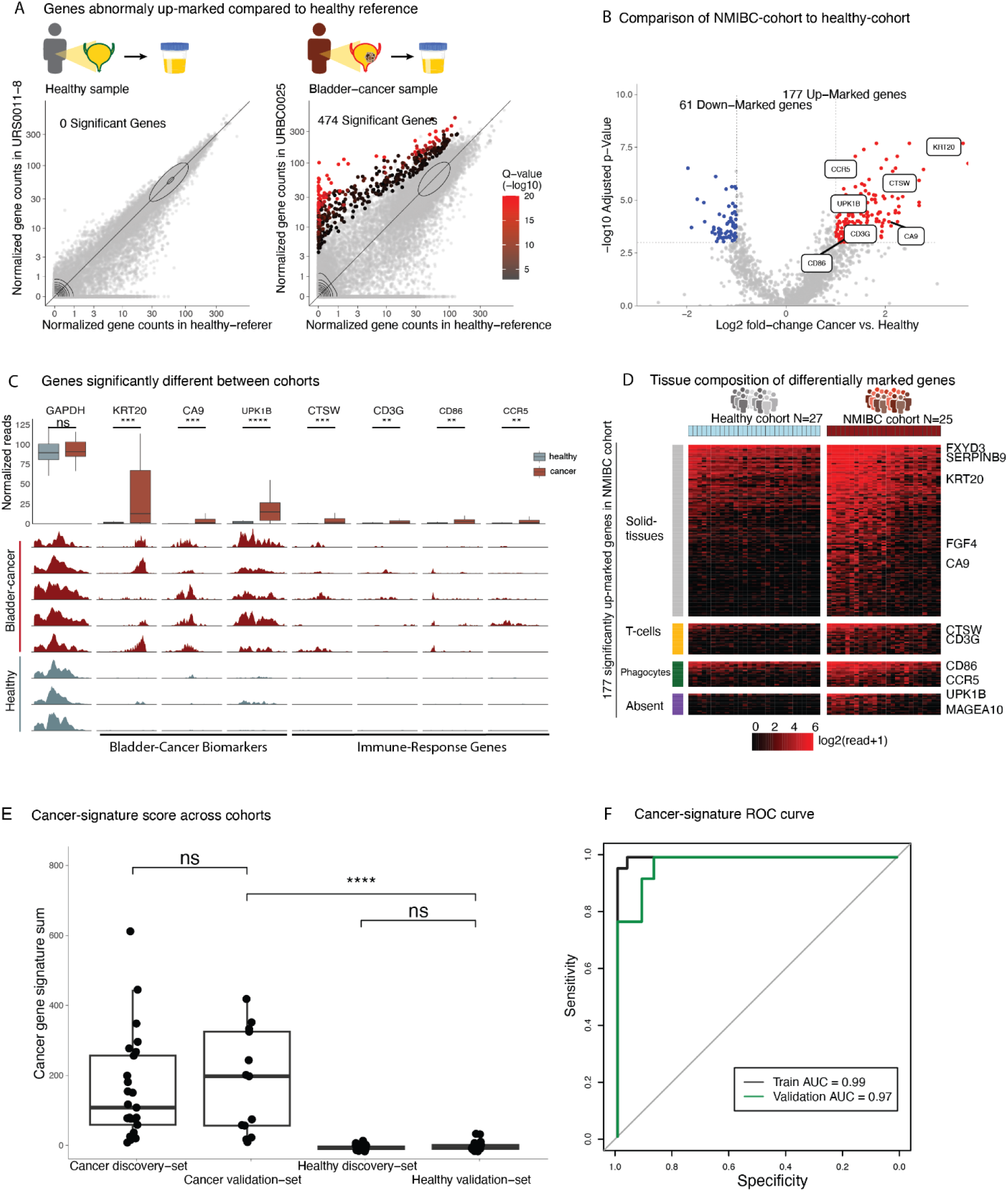
Pathology-driven changes in NMIBC cohort. A. Detection of genes with significant high coverage in a sample from a healthy urine sample (left) and the urine of a patient diagnosed with NMIBC (right). For each gene, we compared mean normalized coverage in a reference healthy cohort (methods, x axis) against the normalized coverage in the cfChIP-seq sample (y axis). Significance tests whether the observed number of reads is significantly higher than expected based on the distribution of values in healthy samples (Methods). B. Cohort comparison across genes. Statistical significance (y-axis) and fold change of NMIBC group over healthy (x-axis) of genes in both groups. Significance values represent FDR-corrected q value of likelihood ratio test of two groups (methods). C. Top, normalized count distribution of representative genes significantly different between the two cohorts (red boxes are NMIBC cohort samples, gray boxes are healthy samples). Bottom, gene browser view of representative genes significantly different between the two cohorts (red tracks are NMIBC cohort samples, gray tracks are healthy samples). D. Heat map representation of 177 genes that are significantly higher in the NMIBC cohort compared to the healthy cohort. Rows (genes) are clustered by cell type category. Heat map values (colors) are log transformed normalized values (log2 (1+normalized gene reads)). E. Comparison of cancer signature levels across the healthy and NMIBC discovery cohorts and the validation cohorts. Boxplot description and statistical test as in figure 2D. F. The Receiver Operating Characteristic (ROC) curve illustrates the performance of a classifier using the cancer signature by plotting the true positive rate (sensitivity) on the y-axis against the false positive rate (1-specificity) on the x-axis.

In contrast to the major differences in urine cfChIP-seq, no differences were found between the plasma cfChIP-seq results of healthy individuals and those diagnosed with NMIBC (Fig. S4B-C), reiterating the value of urine as a valuable tool for study and diagnosis of urinary system pathologies. To ensure these differences in signal are not the results of the age differences between cohorts, we sampled three males ages over 70 without bladder cancer and observed little difference in gene signal (6 or less significantly up-regulated promoters) compared to our healthy cohort. These promoters do not overlap with the signal we see in cancer patients (Supplementary Fig. S4D).

### Changes in cfnDNA of NMIBC patients is driven by cancer-related processes

Within the 177 significantly up-regulated genes in the NMIBC cohort, we observe several bladder cancer expression biomarkers (e.g., KRT20 ^59^, GRHL3 ^60^ FXYD3 ^61^, CA9 ^62^, UPK1B ^63^) indicating oncogenic misregulation at the epigenomic level, observable using urine cfnDNA.

To further investigate the underlying sources of up-regulated promoters in NMIBC samples, we compared this geneset to genes upregulated in different cell types within a reference H3K4me3 ChIP-seq cell-type atlas (Methods, Fig. 4D, S4D). We identified many of these up-marked genes as enriched in T cells (22/177 genes, EnrichR ^64^ ARCHS4 Tissues ^65^ *q*<10^-12^) and phagocytes (18/177 genes, EnrichR ARCHS4 Tissues *q*<10^-16^). These genes have a diminished signal in healthy urine cfChIP-seq and are indicative of a differential immune-cell state, presumably as a response to the tumor. Additional up-regulated genes were categorized as having signal in multiple solid tissues (109/177, solid tissue). Several up-regulated genes had absent signal levels in the reference atlas (15/177, absent), suggesting aberrant activation of pathways that are normally repressed in healthy adults (e.g UPK1B ^63^ and MAGEA10 ^66^). Down-marked genes are enriched in the kidney (EnrichR ARCHS4 Tissues *q*<10^-8^) in agreement with lower levels of kidney observed in cancer samples (Supplementary Fig. S4G-H). This is most likely due to increased signal of other contributing tissues such as T-cells, phagocytes and the tumor in NMIBC samples.

To test if these observations generalize to unobserved urine samples, we defined a cancer signature utilizing gene promoters with a signal in cancer urines but are unmarked in healthy urines (43 genes, termed NMIBC-signature, Supplementary Table S3). This signature performs well in identifying the urines of NMIBC patients when tested against a validation cohort of 23 healthy urine samples and 12 NMIBC urine samples withheld from analysis (Figure 4E-F, Receiver Operating Characteristic (ROC) curve AUC of 0.97, Methods), indicating consistency in abnormal cancer-derived cfnDNA.

## Discussion

In this study, we investigated the presence and characteristics of cf-nucleosomes in urine. Using cfChIP-seq, we demonstrated that urine contains significant quantities of intact nucleosomes carrying various histone marks, including H3K4me1, H3K4me2, H3K4me3, H3K27ac, and H3K27me3. Genomic alignment of the nucleosome-bound cell-free DNA (cfnDNA) is consistent with the expected regions of the accompanying histone mark, suggesting that these histone marks are preserved from their cells of origin.

### Tissue sources of cfnDNA

Focusing on the H3K4me3 promoter mark, we delineated tissue contributions to urinary cf-nucleosomes, identifying diverse cell types, including bladder epithelial cells, kidney cells, and leukocytes. This group of originating cells agrees with prior studies with single-cell RNA sequencing and DNA methylation profiling.

Our observations of sex-based differences in cf-nucleosome profiles further enrich the growing literature on sex-based variability in cfDNA. The enriched population of epithelial cells captured in female samples could account for increased amounts of cfDNA described in female urine^67^

When considering the differences between the cell-free nucleosome populations in urine and plasma, results strongly suggest that in healthy donors, plasma cell-free nucleosomes rarely reach the urine. Megakaryocytes contribute much of the observed contribution to cfDNA in plasma ^36,50^ and are not observed in urine cf-nucleosomes, while kidney cf-nucleosomes are absent in plasma cell-free nucleosomes. Additionally, the presence of plasma cf-nucleosomes originating from the Y chromosome in mothers of male fetuses, coupled with their absence in urine, strongly supports this conclusion. This barrier can be breached in various kidney pathologies.

### Urine cfDNA Fragmentation and origin

The observed cfnDNA fragmentation pattern in urine displays characteristics consistent with digestion patterns observed in plasma and of cfDNA in urine, suggesting a two-phase digestion mechanism—initial cleavage during apoptosis or necrosis (as seen in plasma cfDNA) followed by further digestion within the urine (similar to urine cfDNA). We show that plasma circulating cf-nucleosomes does not traverse into urine under healthy conditions. In contrast, evidence of transrenal cfDNA^68^ suggests that unbound plasma circulating cfDNA can pass to the urine. These findings suggest two different sources of cfDNA in urine and the distinction between nucleosomal cfDNA (recent cell death in the urinary tract) and non-nucleosomal cfDNA (either degradation of cfnDNA or transrenal cfDNA). This contrasts with plasma cfDNA, where nucleosomes dominate the circulating cfDNA pool ^16–18^. Taken together, these suggest that non-nucleosomal cfDNA fragments are efficiently removed from the plasma through glomerular filtration (see model).

### Application in pathophysiology

Beyond cell-type variability, we used urine cf-nucleosomes to detect differences in cell-states. In the urine of NMIBC-patients we observed increased signal of bladder-cancer expression markers and genes indicative of an immune response. These results situate urine cf-nucleosomes as a powerful biomarker for disease detection and monitoring. Longitudinal studies utilizing cf-nucleosomes from urine can further connect these findings to disease recurrence and treatment efficacy based on modulations in cancer- and immune-derived cf-nucleosomes.

### Further research

Some open questions remain regarding urine circulating cf-nucleosomes: Unlike plasma, urine is not homeostatically maintained, making it a highly variable environment. External factors like nutrition, liquid intake, and the female menstrual cycle could affect results in a manner unexplored here. These conditions can also affect the half-life of histone post-translational modifications and cf-nucleosomes. Additionally, all samples in this study were collected from individuals with properly functioning kidneys. Several kidney disorders are characterized by increased permeability of the glomeruli (proteinuria), which could lead to plasma cf-nucleosomes leaking into urine. Finally, while the H3K4me3 promoter mark is effective in cell-type and state annotation, additional information can be obtained using additional histone marks, such as H3K27ac for the annotation of active enhancers and the H3K36me3 mark for active transcription. We have only explored a small subset of the diverse histone-marks, with many more holding information on other cellular processes.

## Conclusion

This study positions urinary cf-nucleosomes as a potent tool for the study of human physiology and disease. By situating these findings within the broader landscape of cfDNA and epigenetic research, we demonstrate the unique advantages of these analytes in studying localized and systemic processes. Histone marks’ dynamic and tissue-specific insights promise to advance basic research and clinical applications, paving the way for more precise and comprehensive biomarker discovery.

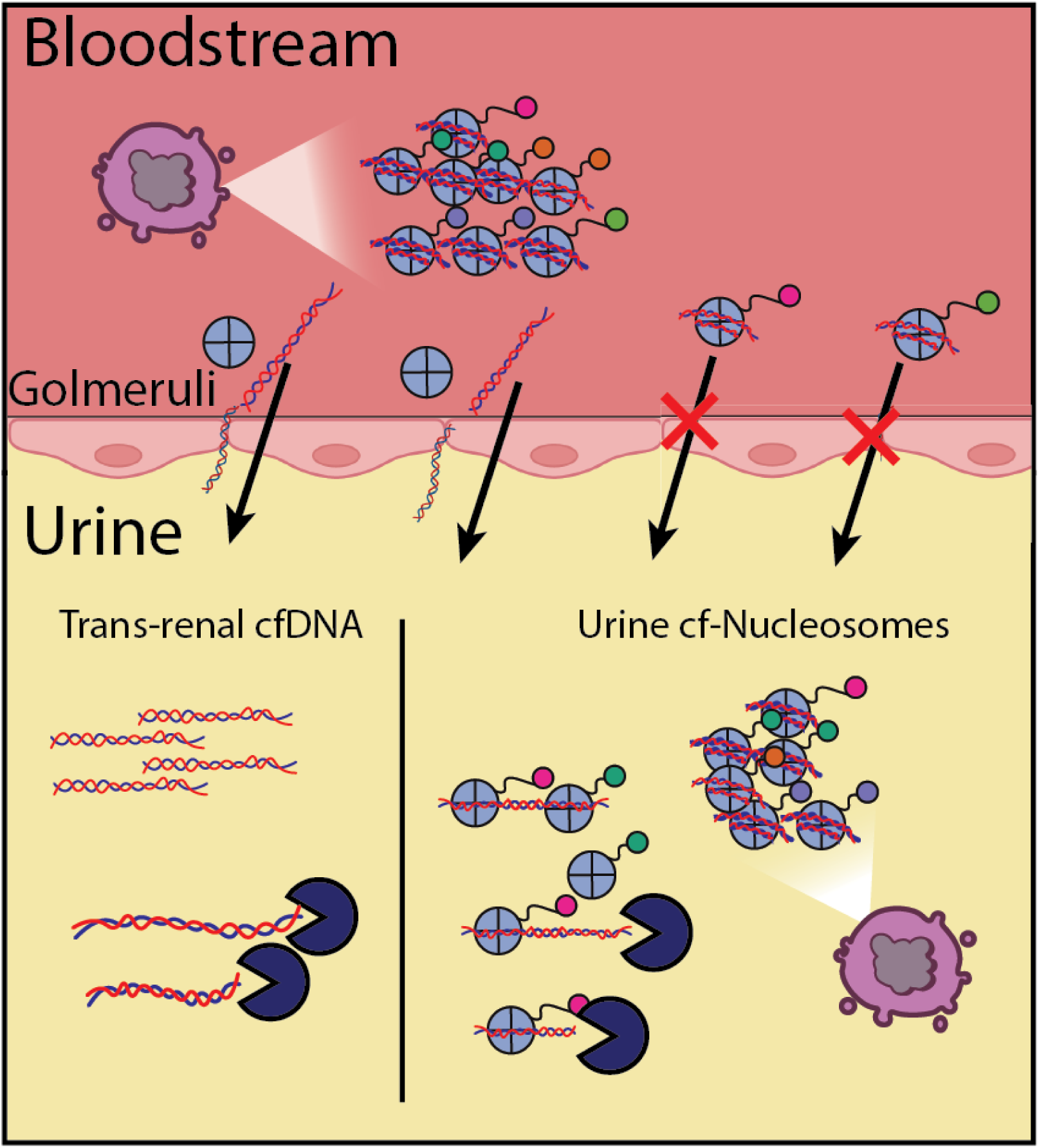

## Model of source of urine cfDNA

Dying cells release cf-nucleosomes in the bloodstream that cannot traverse glomerular filtration into urine, as opposed to unbound-cfDNA which is termed trans-renal cfDNA after passage into urine. Transrenal cfDNA is rapidly digested in urine due to hyperactive DNASE I.

Cells and tissues with direct contact to urine (e.g Kidney, Bladder epithelium and some immune cells) contribute modified cf-nucleosomes to urine after cell death which are also rapidly digested similarly to trans-renal cfDNA, and contribute to the general urine cfDNA pool.

## Materials and Methods

### Donors

All clinical studies were approved by the relevant local ethics committees. The study was approved by the Ethics Committees of the Hadassah-Hebrew University Medical Center of Jerusalem. I Informed consent was obtained from all individuals before blood and urine sampling.

### Sample collection

∼20 ml urine samples were collected in standard plastic collection cups supplemented with EDTA to a final concentration for 10mM to prevent DNase digestion. Samples were transferred to cone 50ml tubes and centrifuged immediately at room temperature at a speed of 3000g for 10 minutes. Urine supernatant was transferred to a second 50 ml cone tube whilst being careful not to disturb any cell debris. The urine supernatant was centrifuged and transferred again using the same settings as before to ensure no cellular debris. The doubly centrifuged urine was used for the cfChIP-seq experiments. Urine cfChIP-seq was performed immediately or after storage at 4C for at most 2 weeks per our preanalytic experiment results (see Supplementary Note).

Plasma samples were collected as described in Sadeh et al. 2021. Briefly, Blood samples were collected in VACUETTE K3 EDTA tubes and transferred to ice. The blood was centrifuged (10 min, 1,500g, 4 °C) and the supernatant was transferred to a fresh 15-ml tube and centrifuged again (10 min, 3,000g, 4 °C). The supernatant plasma was used for ChIP experiments. The plasma was used fresh or flash-frozen and stored at −80 °C for long-term storage.

### Immunoprecipitation, NGS library preparation, and sequencing

Immunoprecipitation, library preparation, and sequencing were performed by Senseera LTD as previously reported in Sadeh et al. ^36^ with slight modifications. Briefly, ChIP antibodies were covalently immobilized to paramagnetic beads and incubated with urine at room temperature overnight or plasma at 4C overnight. For paired-end Illumina sequencing using the NextSeq 550 platform, barcoded sequencing adaptors were ligated to chromatin fragments and DNA was isolated according to the manufacturer’s protocol.

### Sequencing analysis

Paired-end reads were processed by demultiplexing and aligned to the human genome (hg19) using bowtie2 (2.4.2) with ‘no-mixed’ and ‘no-discordant’ flags. Fragments with low alignment scores (flag -q 2) and duplicates were discarded. See Supplementary Table 1 for total number of reads, alignment statistics and number of unique fragments for each sample. BAM files were processed by samtools (1.7) to BED files and by R (4.0.2) scripts (see ‘Code availability’).

Preprocessing of sequencing data was performed as previously described in Sadeh et al. ^36^. Briefly, we constructed a TSS catalog using genomic annotations from ChromHMM, Roadmap Epigenomics, ENSEMBL and UCSC Genome Browser. The human genome was segmented into discrete windows representing gene TSS and background. The sequenced reads covering each of these regions were summed with non-specific reads estimated per sample and then subtracted - resulting in the specific signal in every window. Signal -counts were normalized and scaled to 1 million reads in healthy reference accounting for sequencing depth differences. Detailed information regarding these steps can be found at supplementary note in Sadeh et al.^36^

### Epigenomics atlas

We downloaded 414 samples of H3K4me3 ChIP-seq aligned read data from the ENCODE Epigenomics consortium (https://www.encodeproject.org/) and BLUEPRINT database (https://www.blueprint-epigenome.eu/). BAM files aligned to the hg19 human genome version were processed into BED format exactly as described above. Hg38 aligned BAM files were first converted into hg19 and then processed as above.

List of all cell types used as reference data is in Supplementary Table S5. The BED files were then processed with the same scripts as cfChIP-seq samples.

### H3K4me3 ChIP-seq of urine exfoliated cells

Urine samples were collected as above with the exception of urine exfoliated-cells collection after initial centrifugation. After moving urine to a fresh tube, pelleted cells were washed using ice-cold 1X PBS supplemented with protease inhibitors (Roche cat. 5056489001). Cells were counted [Bio-Rad TC20 Cell Counter] and resuspended using supplemented NP buffer [10 mM Tris pH 7.4, 50 mM NaCl, 5 mM MgCl2, 1 mM CaCl2, and 0.075% NP-40, supplement with fresh 1 mM β-mercaptoethanol, 500 μM spermidine]. Chromatin was digested with Micrococcal nuclease [Worthington Biochemical LS004798] at 5 units per 1 million cells. Chromatin was collected by centrifugation (14,000xg, 10 minutes at 4C) and quantified (Qubit 4 Fluorometer, Agilent TapeStation). Native H3K4me3 ChIP-seq was performed in 500ul SDSless RIPA [10 mM Tris pH 8.0, 140 mM NaCl, 1 mM EDTA, 1% Triton X-100] or Bovine Serum Heat inactivated (Rockland D500-05-0500) using the same protocol as for urine cfChIP-seq. Data processing was also done as above.

### Urine and plasma cfDNA sequencing

cfDNA from urine and plasma samples (collected as above) was extracted using Zymo cfDNA extraction kit [Zymo Research ZR-R1072] according to manufacturer’s instructions. Sequencing libraries were prepared as above. Data processing was also done as above.

### Tissue signatures

Using the ENCODE and Roadmap Epigenomics metadata tables we defined sets of samples that belong to a tissue or group of tissues (see Supplementary Table 12). We then defined for each group of samples the set of specific windows as meeting the following criteria: 1. The window is not predefined as a background region in the genome 2. The window is not located on the Y-chromosome 3. At least two samples of the window satisfy the condition that they have >10 normalized reads, and <5 in all other samples outside the group, or >30 normalized reads and <10 in all other samples - the last condition allows minimal signal from contaminating populations such as immune-populations in solid tissues. Groups for which we found less than 20 specific windows were considered to be without signature. The signature-scores are defined as the sum of normalized reads across signature windows divided by window lengths (normalized reads / kb).

### Cancer-signature (Figure 4E-F)

Significantly differentially up-marked genes in the NMIBC-cohort were filtered for those with a maximum of 3 normalized reads in healthy samples. This set of 43 genes was termed cancer-signature (Table S3).

### Statistical analysis

All custom statistical analyses are thoroughly described in Sadeh et al.^36^ and briefly described below. Computed p-values were corrected for multiple hypotheses using FDR and estimated as q values (R function p.adjust()).

**Comparison to background** (Figure 2B) - We test whether the total sum of reads over a collection of windows (a signature, gene promoter windows, etc.) is larger than we would expect from the background signal. The null hypothesis is that the number of reads in the windows of interest is Poisson distributed according to the estimated background rate at these windows.

**Comparison to healthy reference** (Figure 4A, S4B) - We test whether the total sum of reads over a collection of windows is higher than we would expect according to mean and variance in healthy donor reference (either in plasma or in urine references).

**Comparison of two groups of samples** (Figures 4B, S4C) - For each gene in each group, mean and variance were estimated under a model distribution. We use a likelihood ratio test (LRT) where, under the null assumption, the distribution of the statistic is approximately a Chi-squared distribution with 2 degrees of freedom, which allows us to compute the p-value.

## Acknowledgments

We thank Oliver Rando, Eithan Galun, Tommy Kaplan, Ronen Sadeh, and the members of the Friedman lab for discussions and comments on this manuscript. This work was supported by the European Research Council’s AdG Grant cfChIP 101019560 (to N.F.) and Israel Science Foundation IPMP Grant 3751/21 (to N.F).

## Author contributions

Conceptualization: ML,IS,NF

Methodology: ML,IS

Sample Acquisition: BA

Investigation: ML

Visualization: ML

Funding acquisition: NF

Project administration: ML,IS,TFM,NF

Supervision: NF, TFM

Writing – original draft: ML

Writing – review & editing: ML,IS,BA,TFM,NF

## Data availability

All datasets used in this study are in the process of being deposited to public repositories.

## Code availability

All script files used in the analysis in this manuscript will be available upon request. R code for processing cfChIP-seq data is available at https://github.com/nirfriedman/cfChIP-seq.git, set-up files for analysis are available in the Zenodo repository: https://doi.org/10.5281/zenodo.3967253.

## Supplementary information

- Table S1 - Sequencing statistics for samples sequenced in this study
- Table S2 - Bladder cancer patients clinical information
- Table S3 - Bladder cancer signature genes
- Table S4 - List of all cell-types used as reference data from published ChIP-seq datasets
- Fig. S1-S4
- Supplementary Note

---

**Figure S1:**
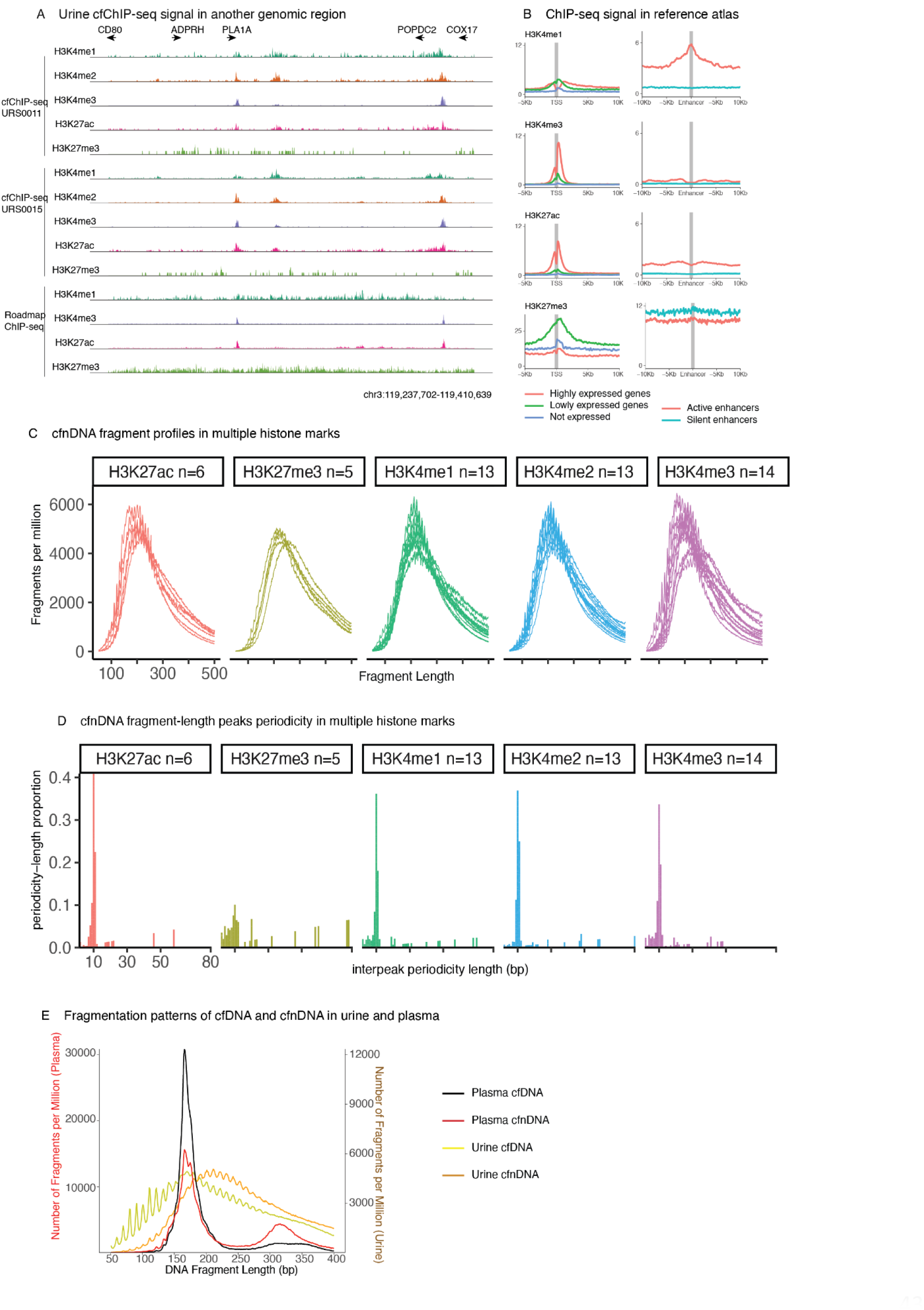
Urine cfChIP-seq fragment size distribution and periodicity. A. Genome browser view as in Fig. 1B B. DNA fragment size distribution across 5 histone PTMs: H3K27ac (N=6), H3K27me3, (N=5), H3K4me1 (N=13), H3K4me2 (N = 13),H3K4me3 (N=14). Each row depicts the number of fragments (opacity) for each fragment length (x axis). Colors indicate different modifications. C. cfnDNA fragment distributions per histone mark. X axis presents DNA fragment length, y axis are urine cfChIP-seq samples and colors represent several histone modifications with shade representing the number of fragments per million. D. Proportion of interpeak periodicity length in each of the histone modifications in this study. In each histone PTM, the length between local maxima in DNA fragments is tallied for each sample, the proportions are then plotted. E. Fragmentation pattern from urine ChIP-seq cfnDNA, urine cfDNA (brown and orange lines) and from plasma cfChIP-seq cfnDNA and plasma cfDNA (red and dark-red lines).

**Figure S2:**
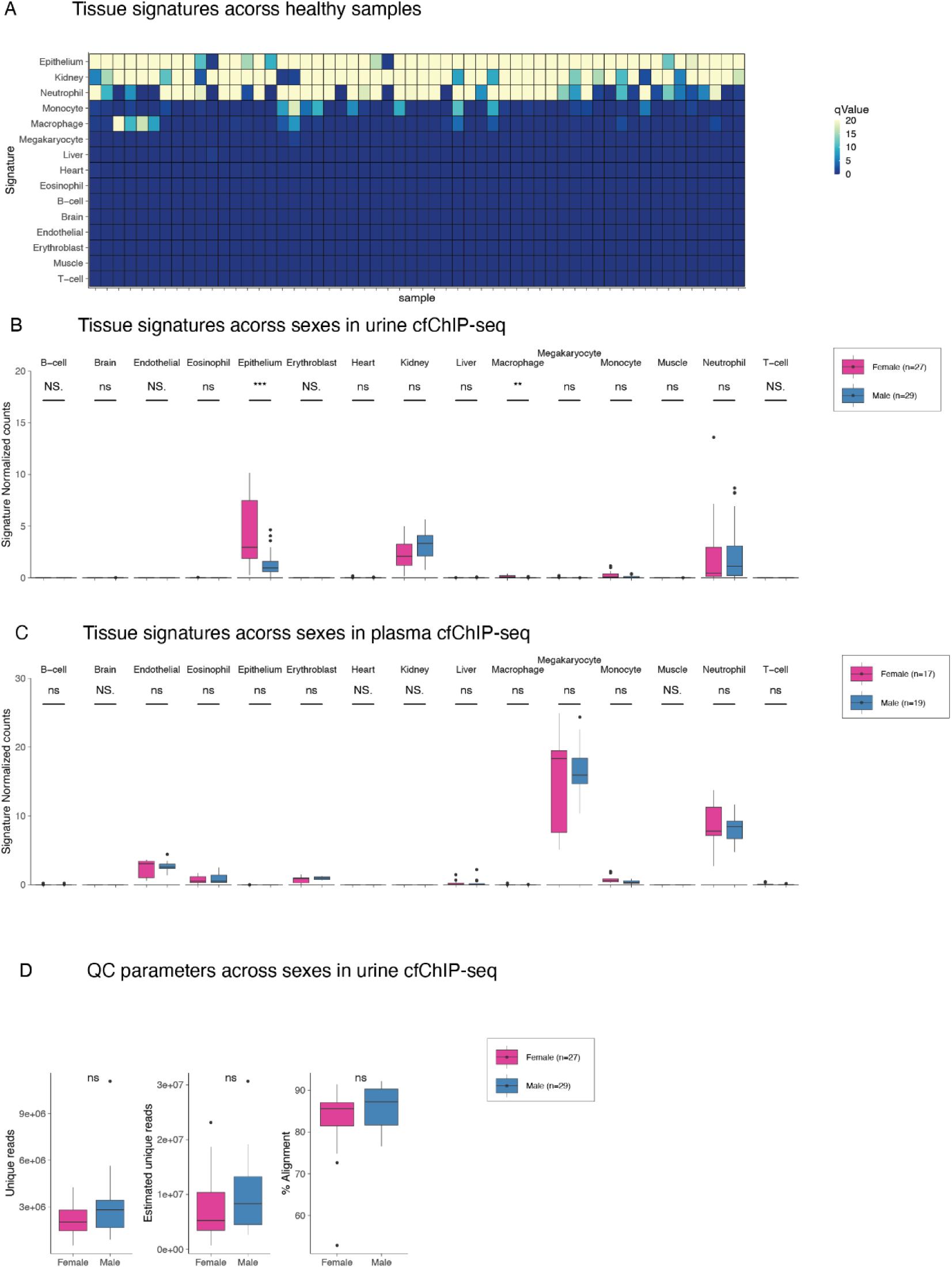
Tissue signatures. A. Tissue signatures (rows) of all healthy samples (columns, N=56) color in heatmap represents the adjusted p-value (q-value) of the null hypothesis that the signature’s signal is equivalent to noise. B. Tissue signature comparison between male and female samples of urine cfChIP-seq as in Fig. 2D. C. Tissue signature comparison between male and female samples of plasma cfChIP-seq as in Fig. 2D. D. Quality control (QC) parameters in both sexes. Statistical comparison done via Mann-Whittney U-test, “ns” - “not significant”

**Figure S3:**
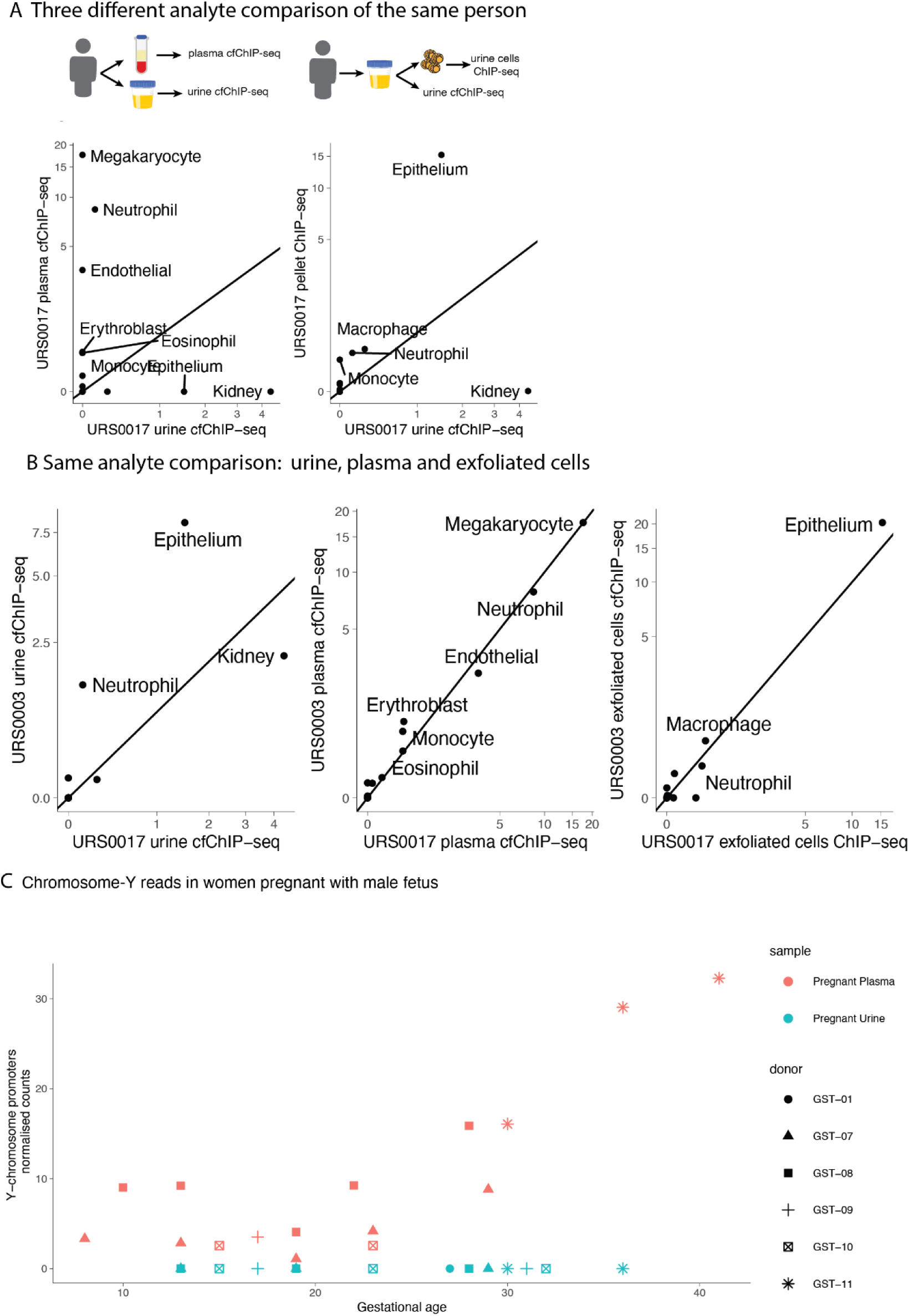
Comparison of Urine, exfoliated urine-cells and plasma. A. Same as Figure 3A - signature comparison of three analytes from the same individual. B. signature comparison of the same analyte from two different individuals. Left, urine cfChIP-seq. Middle, plasma cfChIP-seq. Right, urine exfoliated cells ChIP-seq. C. Normalized promoter coverage in the Y-chromosome in urine and plasma samples from females pregnant with male fetuses, across different gestational ages by week. data-points are colored by sample type and shapes indicate the pregnant donor.

**Figure S4:**
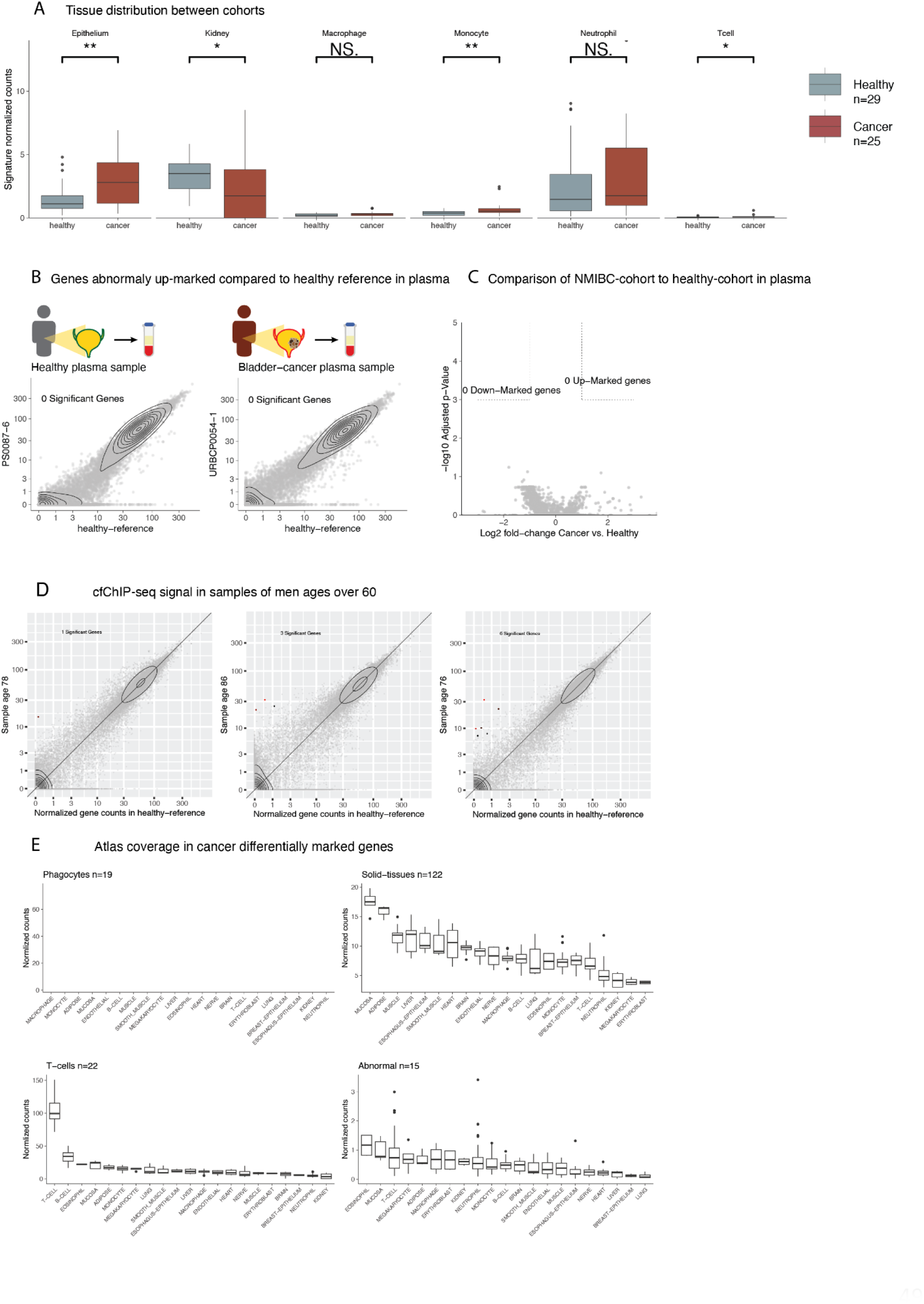
Urine cfChIP-seq of NMIBC cohort. A. Tissue distribution of the healthy (gray boxes) and NMBC (dark red boxes) cohorts. Boxplot descriptions and statistical tests as in figure 2D. B. Detection of genes with significant high coverage in a sample from a healthy plasma sample (left) and the plasma of a patient diagnosed with NMIBC (right). For each gene, we compared mean normalized coverage in a reference healthy cohort (x axis) against the normalized coverage in the cfChIP-seq sample (y axis). Significance tests whether the observed number of reads is significantly higher than expected based on the distribution of values in healthy samples (Methods). C. Cohort comparison across genes. Statistical significance (y-axis) and fold change of NMIBC group over healthy (x-axis) of genes in both groups. Significance values represent FDR-corrected q value of likelihood ratio test of two groups (methods). D. Comparisons of gene signals from urine samples from male donors (y-axis) ages 76 (left) 86 (middle) and 76 (right) to a healthy cohort of urine samples (x-axis). For each gene, we compared mean normalized coverage in a reference healthy cohort (methods, x axis) against the normalized coverage in the cfChIP-seq sample (y axis). Significance tests whether the observed number of reads is significantly higher than expected based on the distribution of values in healthy samples (Methods). E. 177 significantly up-marked genes in the NMIBC cohort are categorized by tissues which they are highly marked in (methods). Boxes are different tissues in the ChIP-seq reference atlas (x-axis), box height is the mean normalized gene count for the set of genes in each category.

## Supplementary Note

Assessment of the contribution of cf-nucleosomal DNA to total urine cfDNA

We use the H3K4me3 marked cf-nucleosomes in urine for the following estimate:

- The human (diploid) genome is 6.4 billion bp
- The human diploid genome weighs 6.4 pg
- Nucleosome length + linker is approx. 200 bp
- 30 million nucleosomes in a diploid cell
- of these 0.5-1% are marked by H3K4me3 ^2^ = 150,000 K4me3 nucs per cell.
- in urine H3K4me3 cfChIP-seq we estimate the library complexity to be ∼10M unique reads per sample (2ml urine)
- This is the equivalent of 66 diploid genome equivalents (10e6 estimated nucs per sample /1.5e5 nucs in a single cell)
- The human diploid genome weighs 6.4 pg ^1^
- 6.4pg * 66 GE = 0.42 ng from 2 ml urine sample per cfChIP-seq = 0.21 ng/ml

Below are estimates of total urine cfDNA concentrations in 8 studies ^3–10^

Evidently the reported concentrations show high variability ranging from an average of >1 up to 30 ng/ml.

This can be the results of the different extraction kits used as well as sample handling and storage as we show in our study pre-analytics.

Our estimate of 0.21 ng/ml of nucleosomal cf-dna in urine positions their contribution within an order of magnitude of other estimates presented, indicating it is a non-trivial contribution to urine cfDNA pool.

(Supplementary Note fig. 1)

**Figure 1.**
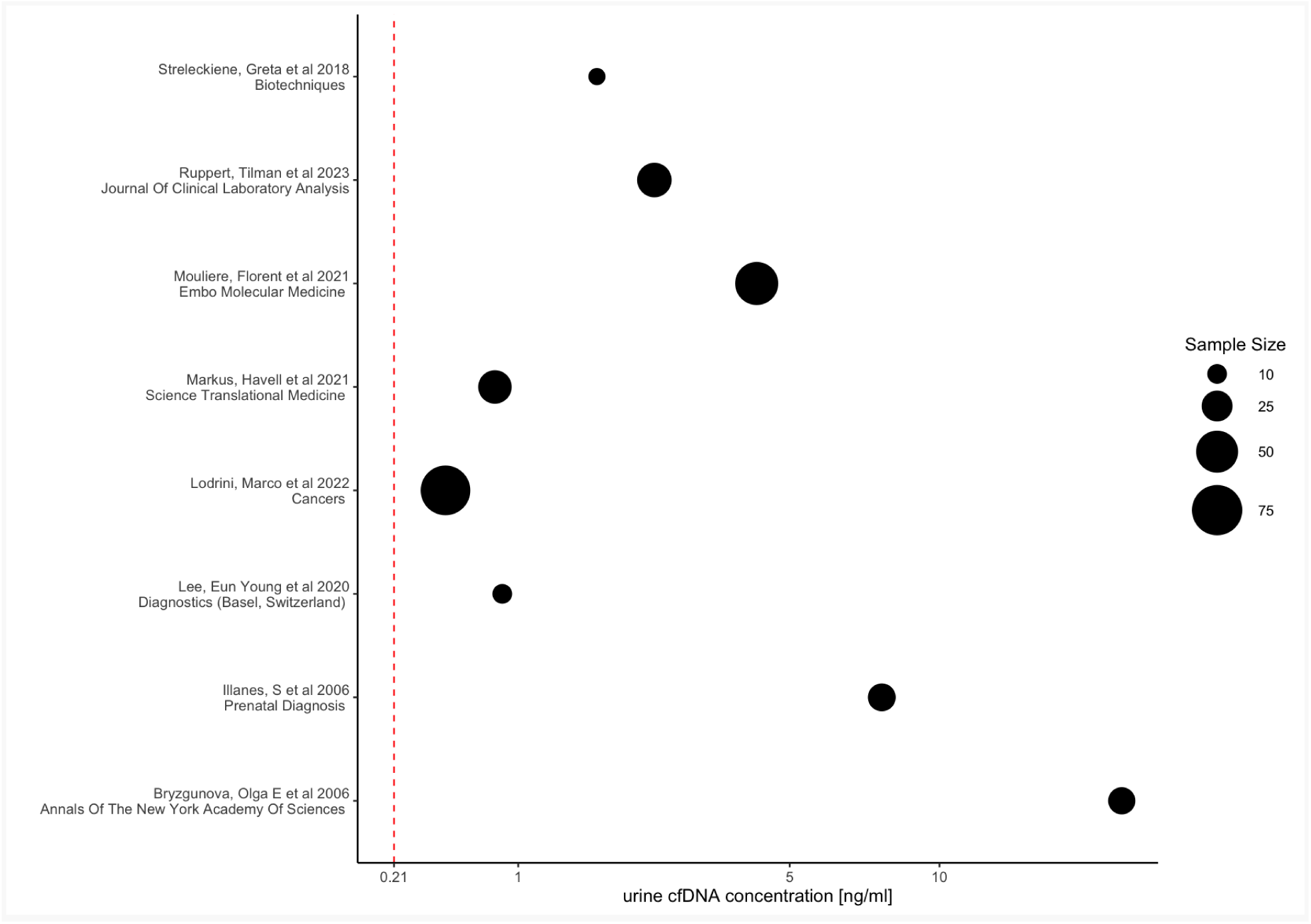
Urine cfDNA concentrations in previous publications. Urine cfDNA concentrations (x-axis) from 9 recent publications (y-axis) are reported in ng/ml as either point estimation of the mean of samples collected (dot) or as range (line). Red-dashed line represents the contribution of cfDNA from urine cf-nucleosomes as estimated from cfChIP-seq.

## Urine Sample preanalytics

Urine samples collected as specified in the Methods section were kept in three storage conditions and then cfChIP-seq was performed.

Conditions tested were three time points (3 days, 7 days and 14 days) in room temperature or cooling (4C) and three time points (7 days, 14 days and 30 days) in freezing conditions (-80C). The QC parameters observed were the estimated number of unique fragments in the sample, the percentage of unique reads mapping to promoters (% signal in TSS) and the overall percentage of aligned reads. We found urine cfChIP-seq quality rapidly deteriorates in room-temperature but is maintained in 4C and -80C over-time relative to freshly processed samples (Supplementary Note fig. 2).

**Figure 2.**
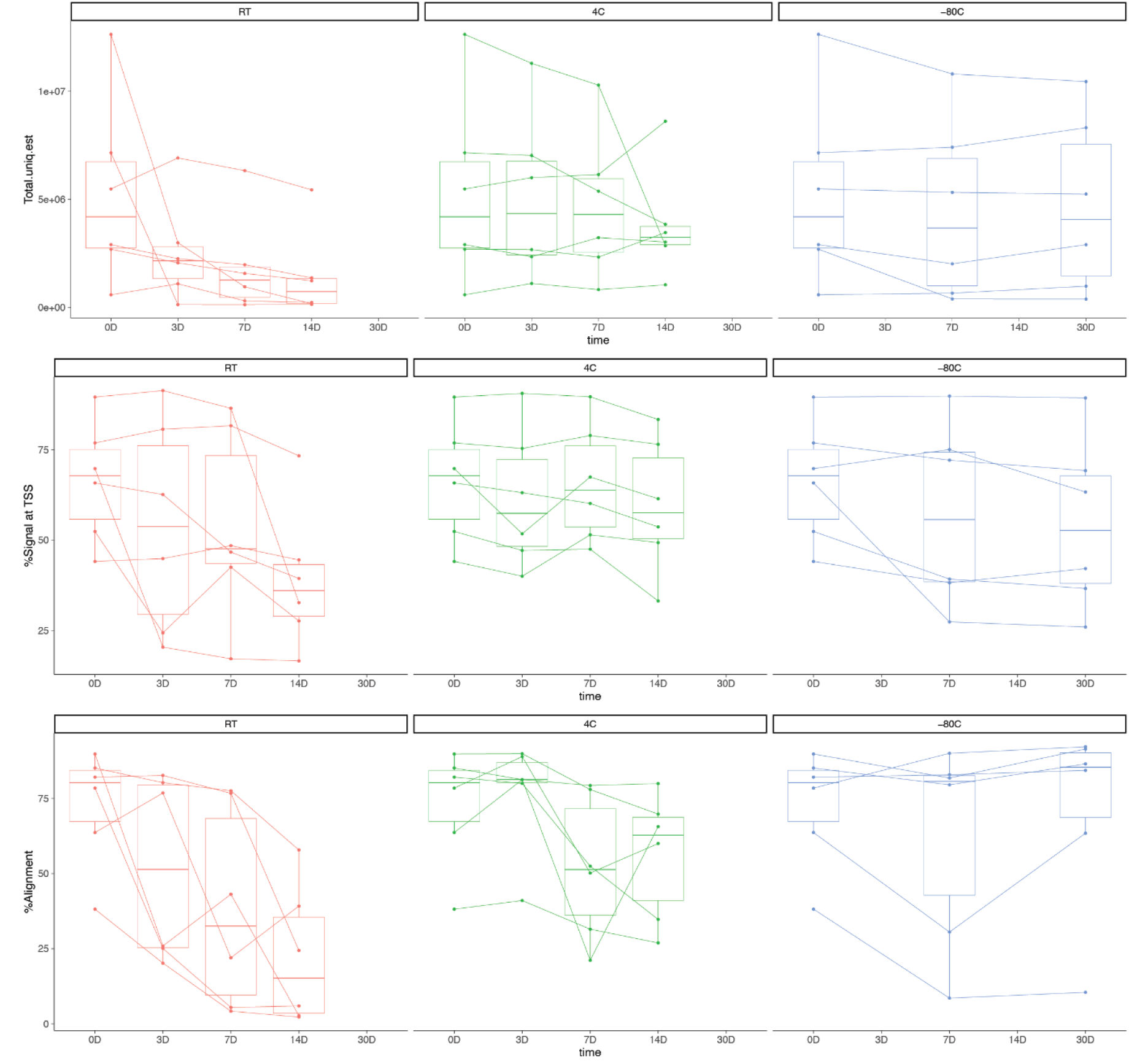
Urine Sample preanalytics: Urine cfChIP-seq was applied to 6 samples (3 Male and 3 Female) after storage in three common lab conditions: room-temperature, 4C and -80C. Samples were processed after 3,7 and 14 days in room temperature and 4C, and after a week and after a month in -80C storage. **Top row,** total unique estimated reads which reflects sequencing library complexity. **Middle row**, %signal in TSS which reflects the proportion of reads that were mapped to promoter windows in the promoter-mark H3K4me3 in urine cfChIP-seq. **Bottom row**, the proportion of aligned reads in sequencing.

